# Oncogenic RAS-MET signal interactions are modulated by P53 status in NF1-related MPNSTs

**DOI:** 10.1101/199026

**Authors:** Matthew G. Pridgeon, Elizabeth A. Tovar, Curt J. Essenburg, Zachary Madaj, Elissa A. Boguslawski, Patrick S. Dischinger, Jacqueline D. Peacock, Flavio Maina, Rosanna Dono, Mary E. Winn, Carrie R. Graveel, Matthew R. Steensma

## Abstract

We previously reported that cooperative RAS-MET signaling drives disease progression in NF1-related MPNSTs, and that MET inhibition results in downstream inhibition of RAS/MAPK in the context of *MET* amplification. This study revealed that response to MET inhibition appeared to be modulated by *P53* gene status. It is currently unclear how P53 function affects kinome signaling and response to kinase inhibition. Here we utilized genetically engineered mouse models with variable levels of *Met* and *Hgf* amplification and differential *p53* status (*NF1*^*fl/KO*^*;lox-stop-loxMET*^*tg/+*^*;Plp-creERT*^*tg/+*^; *NF1*^*+/KO*^*;p53*^*R172H*^*;Plp-creERT*^*tg/+*^; and *NF1*^*+/KO*^*;Plp-creERT*^*tg/+t*^). These NF1-MPNST models were used to assess a novel MET/MEK (i.e. RAS-MET) inhibition strategy and investigate the adaptive kinome response to MET and MEK inhibition. We demonstrate that combination MET (capmatinib) and MEK (trametinib) inhibition fully suppresses MET, RAS/MAPK, and PI3K/AKT activation in P53 wild type tumors, whereas P53-mutant tumors demonstrated sustained CRAF, BRAF, and AKT activation in the presence of combined MET and MEK inhibition. Interestingly, trametinib therapy alone strongly activates MET signaling in *MET* and *HGF*-amplified tumors regardless of P53 status, an effect that was abrogated by the addition of capmatinib. We conclude that P53 alters RAS-MET signaling interactions that drive therapy resistance in NF1-related MPNSTs.

## Introduction

Cooperative RAS/ERK and HGF/MET signaling promotes cancer progression and therapy resistance, yet the mechanisms that mediate these complex signaling interactions are not well understood. Recent studies support the concept that cooperative RAS-MET signaling is influenced by genomic alterations and feedback within the MET and RAS signaling pathways. Neurofibromatosis type 1 (NF1) is a tumor predisposition syndrome caused by germline mutations in the *NF1* gene which encodes neurofibromin, a GTPase-activating protein that regulates RAS signaling (1, 2). Approximately 8-13% of individuals with NF1 will develop malignant tumors, most commonly Malignant Peripheral Nerve Sheath Tumors (MPNSTs) (3). NF1-related MPNSTs are chemorefractory sarcomas that frequently metastasize and have five-year survival rates ranging from 20-50% (4-8). MET-RAS signaling is implicated in NF1-related MPNSTs. MPNSTs often develop from MET-overexpressing plexiform neurofibromas and commonly exhibit *MET* and *HGF* gene amplifications (9, 10). These genomic alterations occur in the backdrop of putative RAS deregulation from germline loss of *NF1*-mediated tumor suppression, and *NF1* loss-of-heterozygosity events in the MPNST cell of origin. Moreover, MET phosphorylation is observed in at least half of MPNSTs and combined inhibition of MET/VEGFR (cabozantinib) mitigated tumor growth in a preclinical MPNST model (11). Aberrant MET signaling is known to drive malignant progression in a variety of RAS deregulated tumors in humans, as well as therapy resistance (12). MET has also been found to drive resistance to RAS pathway inhibition in several cancers, including melanoma and colorectal cancer (13, 14)

We recently showed that MET overexpression, as a result of *MET* amplification in *NF1*-null Schwann cells, is sufficient for malignant transformation and that *MET* amplified MPNSTs respond exquisitely to targeted MET inhibition (15). We also conducted a longitudinal genomic analysis to identify key genetic events underlying transformation of a human plexiform neurofibroma to MPNST (15). Our results revealed that there is early positive selection for *MET* and *HGF* copy number gain during human MPNST progression that precedes both the accumulation of other oncogene amplifications and additional losses of tumor suppressor genes. In order to validate these findings, we designed a unique mouse model of *MET* amplification in *p53* wild type, *NF1*-null myelinating cells (*NF1*^*fl/KO*^*;lox-stop-loxMET*^*tg/+;*^*Plp-cre ERT*^*tg/+*;^ referred to as **NF1-MET**) (15). NF1-MET mice develop MPNSTs in the absence of additional mutations. A comparison of NF1-MET MPSNTs with tumors derived from a closely related *p53* deficient model (*NF1*^*KO/+*^*;p53*^*R172H*^*;Plp-creERT*^*tg/+*^; referred to as **NF1-P53**), and *Hgf*-amplified P53 wild-type model (*NF1*^*KO/+*^*;Plp-cre ERT*^*tg/+*^; referred to as **NF1**), identified distinct MET, RAS, and PI3K/AKT signaling patterns that strongly implicate RAS-MET signaling as a mode of resistance to kinase inhibitors in MPNSTs.

In addition to *MET* and *HGF* copy number variations, NF1-related MPNSTs also exhibit highly complex genomic structural variations leading to additional genomic gains and losses that appear to be non-random and adaptive for MPNST progression (15-17). Consistent genomic alterations have been identified in MPNSTs, most notably mutations in the *CDKN2A* and *TP53* tumor suppressors (18-20). Recent work confirms that clonal heterogeneity is attributable to NF1/P53 and NF1/CDKN2A deficiency due to the formation of tumorigenic stem cells (21). While it is unclear whether P53 and CDKN2A mutations affect kinase signaling, MET activation has been proposed as a stem cell marker in P53-deficient breast cancer lines. A recent study showed that *Met* amplifications promote a stem cell phenotype in the context of P53-deficiency suggesting that the pleiotropic effects of MET activation support the stem cell niche (22, 23). Other studies present evidence that HGF/MET activation is directly regulated through P53 (24, 25). We test the hypothesis that P53 modulates kinome signaling in NF1-related MPNSTs using a novel therapeutic strategy: combination MEK (trametinib) and MET (capmatinib) inhibition. We employ a panel of genetically engineered mouse models that spontaneously develop NF1-related MPNSTs and examine the effect of *MET* and *HGF* amplifications on tumor growth and tyrosine kinase inhibitor (TKI) response. We show that P53 gene status is associated with differential RAS/ERK, MET, and PI3K/AKT signaling patterns in response to both single agent and combined MET/MEK inhibitor therapy. Interestingly, the primary therapy resistance observed with P53 mutations is overcome with MET inhibition, highlighting the importance of RAS-MET signal interactions in NF1-related MPNSTs. Furthermore, we identify a novel mode of intrinsic therapy resistance to MEK inhibitors through rapid activation of MET and AKT.

## Results

### Evaluating the efficacy of MET and MEK inhibition in MPNST models of *Met* amplification and *p53* loss

Previously, we examined the effects of MET inhibition on tumor growth and receptor tyrosine kinase (RTK) signaling in three genomically distinct MPNST models with the MET inhibitor capmatinib (15). Capmatinib (INC280) is an oral, highly selective, and potent MET inhibitor that is well tolerated and has shown clinical activity in advanced solid tumors (26-28). P53 wild type, NF1-MET MPNST tumors (containing 8 *Met* copies) were uniformly sensitive to capmatinib treatment, however P53 deficient, NF1-P53 tumors (containing 4 *Met* copies) and P53 wild type, NF1 tumors (containing 3 *Hgf* copies) were only partially inhibited and demonstrated significant response heterogeneity with sustained activation of ERK and AKT. These results suggest that MET inhibition is overall effective in reducing MPNST growth in MET-addicted MPNSTs, yet P53 deficiency is associated with relative capmatinib resistance. Moreover these results and other TKI studies suggest targeted inhibition of multiple kinase nodes may be required to minimize response variability and abrogate bypass mechanisms of resistance (29-31).

Since RAS deregulation is a hallmark of NF1-related tumors, we chose to evaluate the efficacy of MEK inhibition. The recent clinical success of MEK inhibition (i.e. selumetinib) in NF1-related plexiform neurofibromas highlights the therapeutic potential of targeting MEK in NF1-related peripheral nerve tumors such as MPNSTs (32). In this study, we used the MEK inhibitor trametinib (Mekinist) which is a reversible, highly selective, allosteric inhibitor of MEK1 and MEK2, and is FDA approved for metastatic melanoma. One of the challenges of targeting the MEK/ERK pathway is achieving high level MEK inhibition without systemic toxicity. Trametinib has a strong pharmacokinetic profile with exceptional potency and specificity, oral bioavailability, and long half-life with a shallow C_max_ (peak concentration) to C_trough_ (trough concentration) profile (33). To compare the effects of MET and MEK inhibition in MPNSTs, we performed *in vivo* and *in vitro* preclinical studies with single agent capmatinib or trametinib. We also included the standard chemotherapy agent doxorubicin which is commonly used to treat MPNST patients yet achieves minimal response. Tumorgrafts were established in immunocompromised mice from primary MPNSTs in the NF1-MET, NF1-P53, and NF1 models as previously described (15). Transplanted tumors were allowed to grow until tumor volume was approximately 150mm^3^ and then animals were randomized and treated for 3 weeks with one of the following treatments: 1) vehicle, 2) doxorubicin, 3) capmatinib, 4) trametinib, 5) trametinib + doxorubicin, 6) capmatinib + doxorubicin, 7) capmatinib + trametinib, or 8) capmatinib + trametinib + doxorubicin.

To evaluate the efficacy of MET and/or MEK inhibition in our distinct MPNST models, we analyzed the effect of monotherapy and combination therapy on tumor growth and response variability in NF1-MET (Figure 1), NF1-P53 (Figure 2), and NF1 (Figure 3) tumorgrafts. To expand our analysis of the kinome response to single kinase vs. combined kinase inhibition, we measured the effects of MET and MEK inhibition on the RAS-MET kinome using both MPNST tumorgrafts and cell lines were derived from the NF1-MET, NF1-P53, and NF1 tumor models (15). Since the majority of tumor tissue is necrotic after 21 days of treatment in ‘responsive’ tumors, we evaluated the signaling activity in tumor tissue after 2 days of treatment when the majority of the tumor is viable and necrosis is absent. Specifically, we examined critical RAS-MET signaling nodes in each of the three MPNST tumor models including phosphorylated MET (pMET; Figure 4), phosphorylated ERK (pERK; Figure 5) and phosphorylated AKT (pAKT; Figure 6). Both AKT and ERK signaling are key downstream signaling effectors of RAS and MET and may be critical mediators of the adaptive kinome response to MET and MEK inhibition. To assess global activity changes in each tumor, we examined MET (Figure 4A), ERK (Figure 5A), and AKT (Figure 6A) activity by immunohistochemical staining at 20X and 100X magnification. To further interrogate the kinome response to MET and MEK inhibition, we treated MPNST cells from each MPNST models with capmatinib and/or trametinib for 2 hours (Figure 4-7).

**Figure 1:**
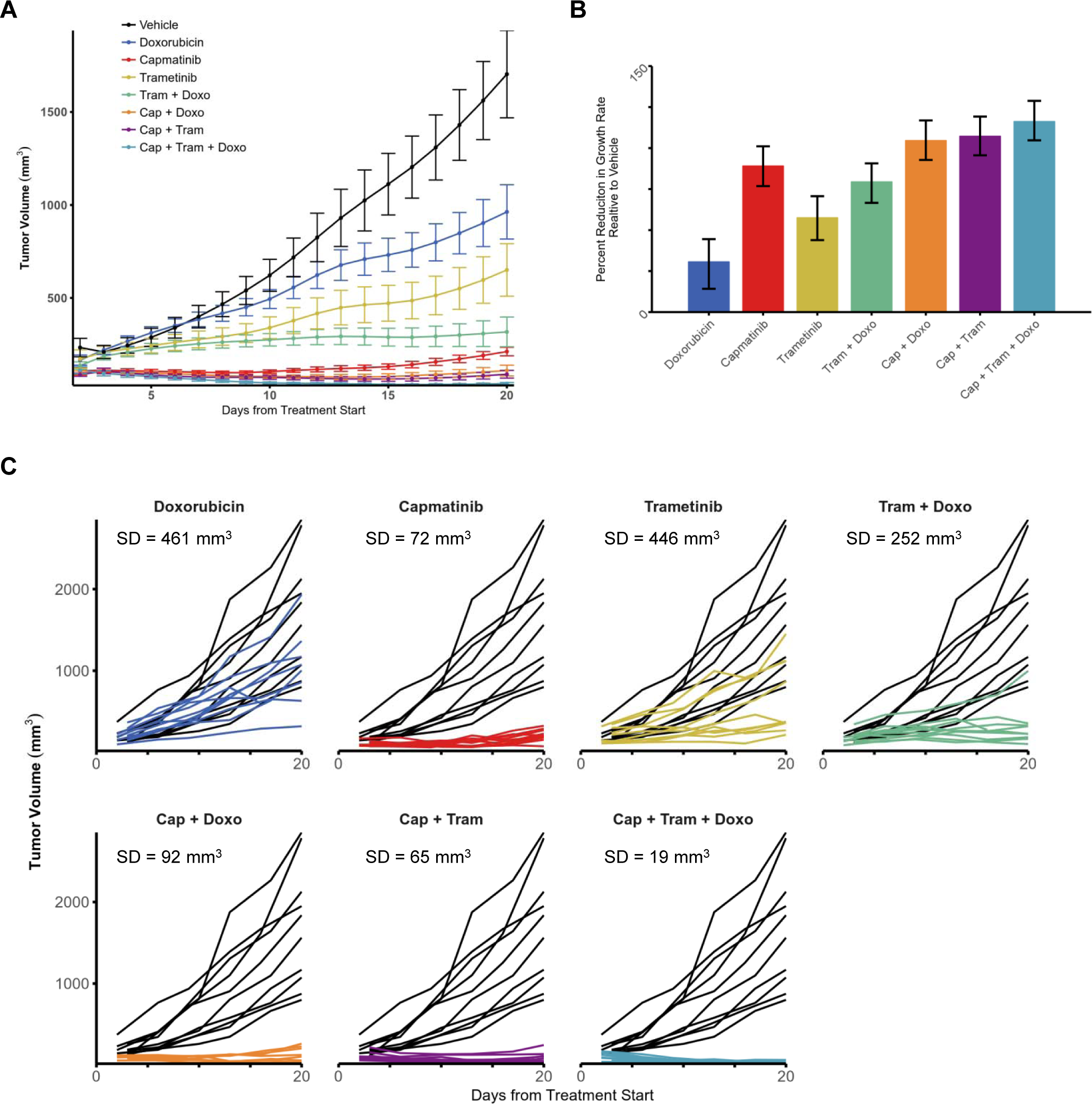
NF1-MET tumorgrafts are exquisitely sensitive to dual kinase MET and MEK inhibition with capmatinib and trametinib. (A) Tumor growth of NF1-MET tumorgrafts is plotted as the spline interpolated means with standard errors. Curves terminate once >70% of mice have been euthanized in the respective treatment group. (B) 95% confidence intervals for the pairwise difference between the growth rates of the select treatments. Statistically significant differences between compared therapies are highlighted in red. (C) Individual tumor growth plotted by treatment (colored lines) compared to vehicle (black lines) over 20 days.

**Figure 2:**
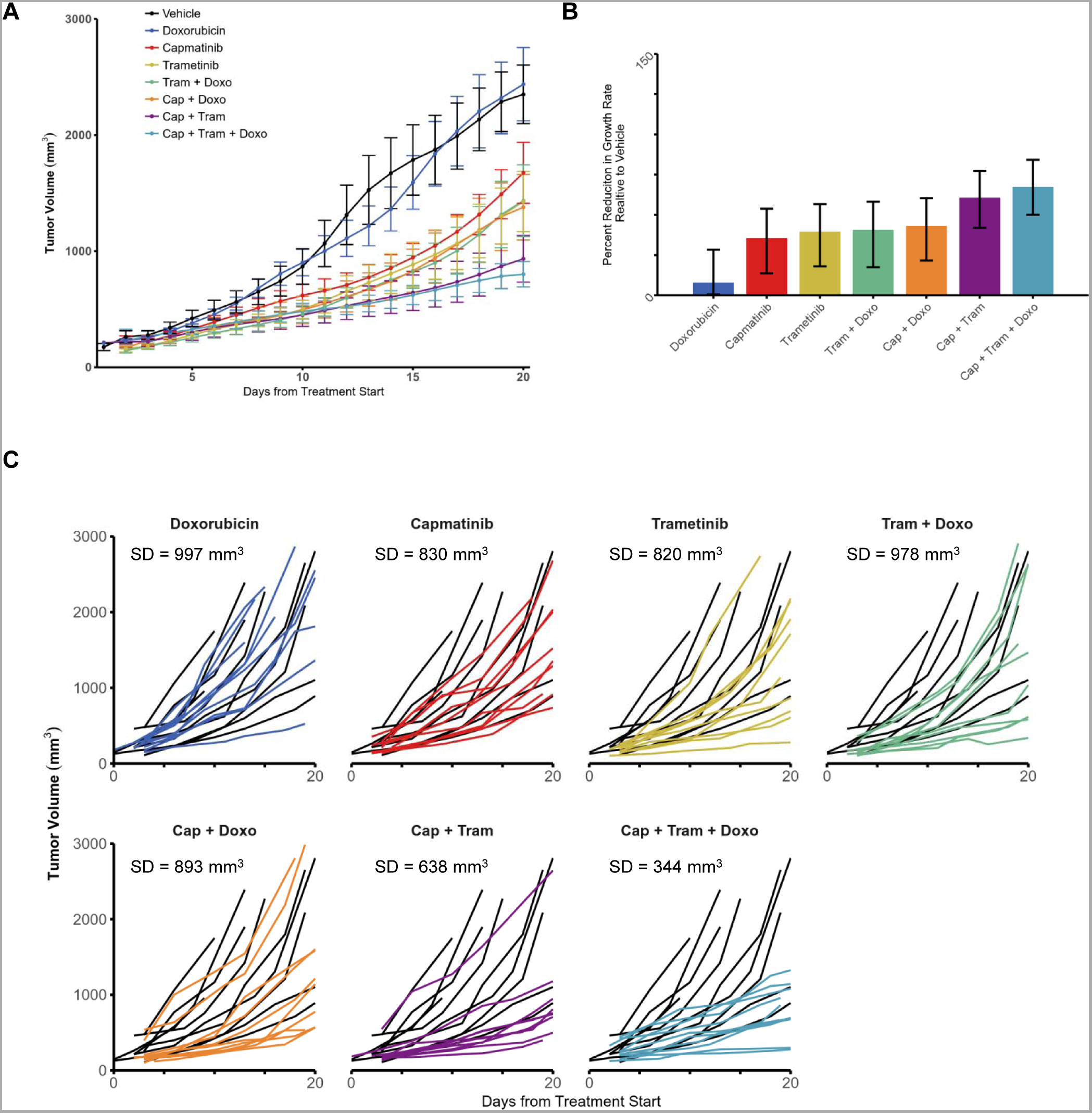
NF1-P53 tumorgrafts grow aggressively and require combined MET and MEK inhibition for substantial growth inhibition. (A) Tumor growth of NF1-P53 tumorgrafts is plotted as the spline interpolated means with standard errors. Curves terminate once >70% of mice have been euthanized in the respective treatment group. (B) 95% confidence intervals for the pairwise difference between the growth rates of the select treatments. Statistically significant differences between compared therapies are highlighted in red. (C) Individual tumor growth plotted by treatment (colored lines) compared to vehicle (black lines) over 20 days.

**Figure 3:**
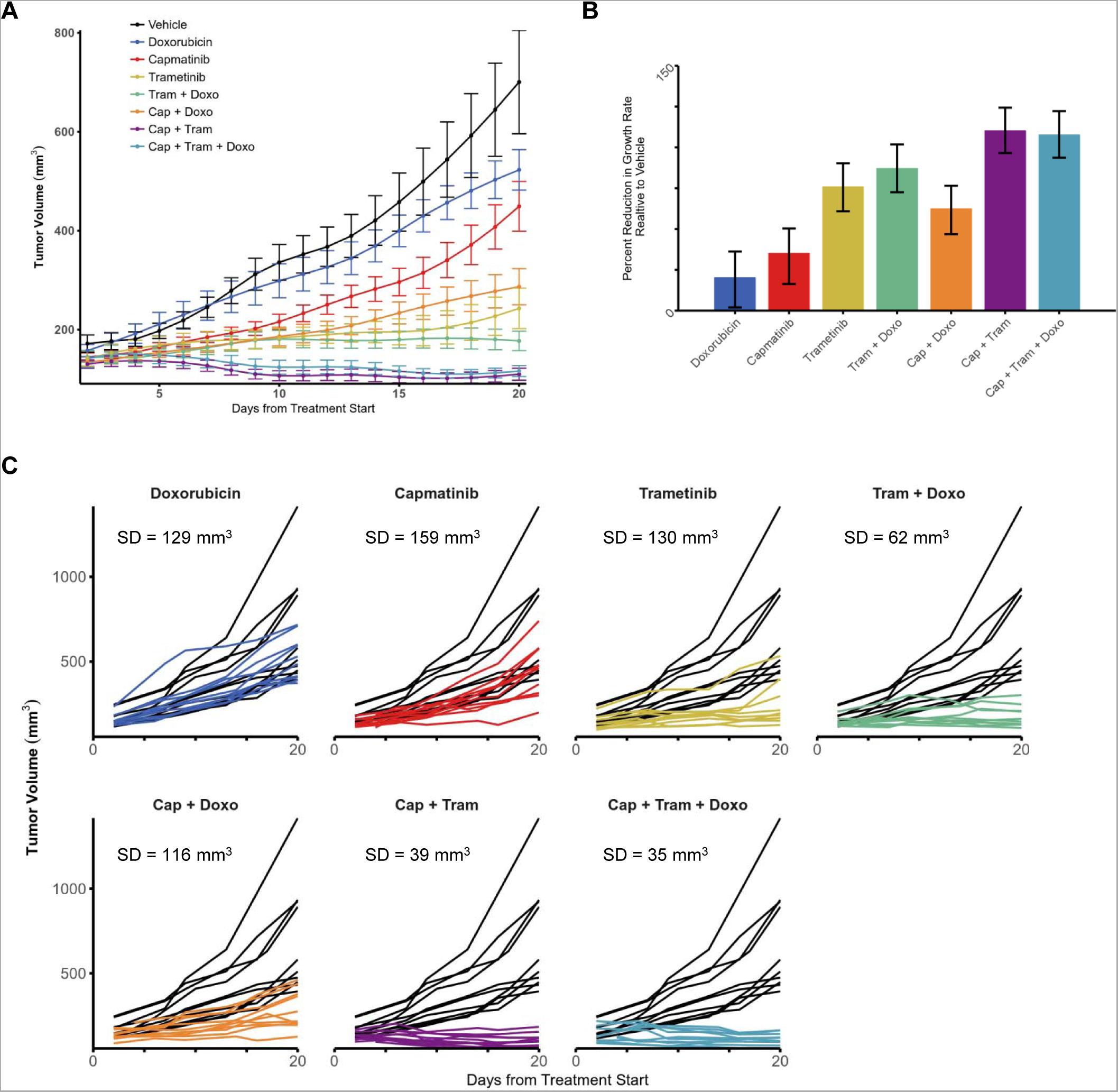
NF1 tumorgrafts are sensitive to dual kinase MET and MEK inhibition with capmatinib and trametinib. (A) Tumor growth of NF1 tumorgrafts is plotted as the spline interpolated means with standard errors. Curves terminate once >70% of mice have been euthanized in the respective treatment group. (B) 95% confidence intervals for the pairwise difference between the growth rates of the select treatments. Statistically significant differences between compared therapies and are highlighted in red. (C) Individual tumor growth plotted by treatment (colored lines) compared to vehicle (black lines) over 20 days.

**Figure 4:**
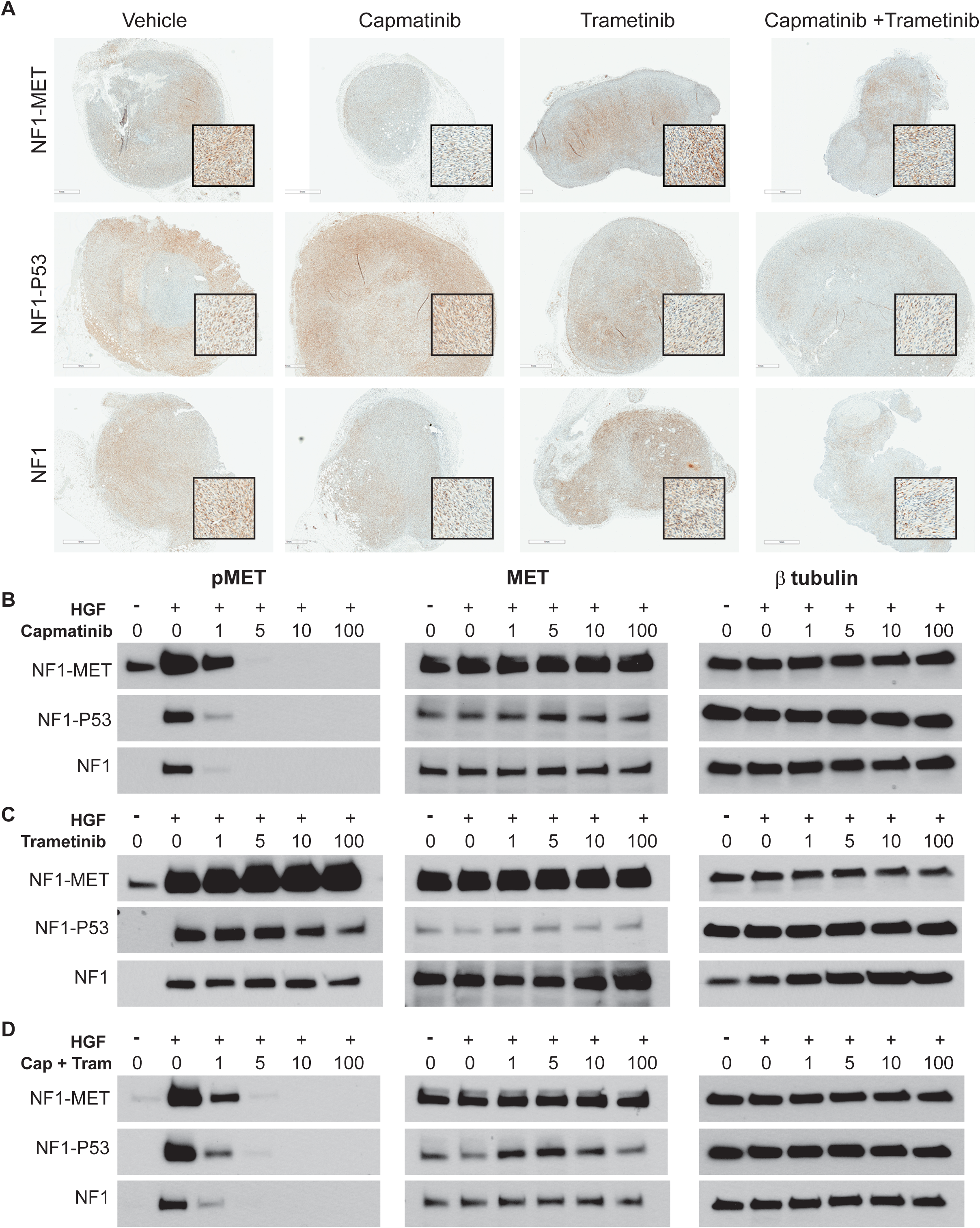
Effects of MET and MEK inhibition on MET activation in NF1 MPNST Models. (A) Immunostaining of pMET (Y1234/1235) in MPNST models that were treated with kinase inhibitors for 2 days (areas of strongest signal are shown in the magnified inset boxes). (B-D) Western Blot analysis of MPNST cells treated with TKIs for 2 hours and +/-HGF for pMET (left panels), total MET (middle panels), and β-tubulin control (right panels). *The total MET exposure for the NF1-MET cells was half the time of the NF1-P53 and NF1 cells.

**Figure 5:**
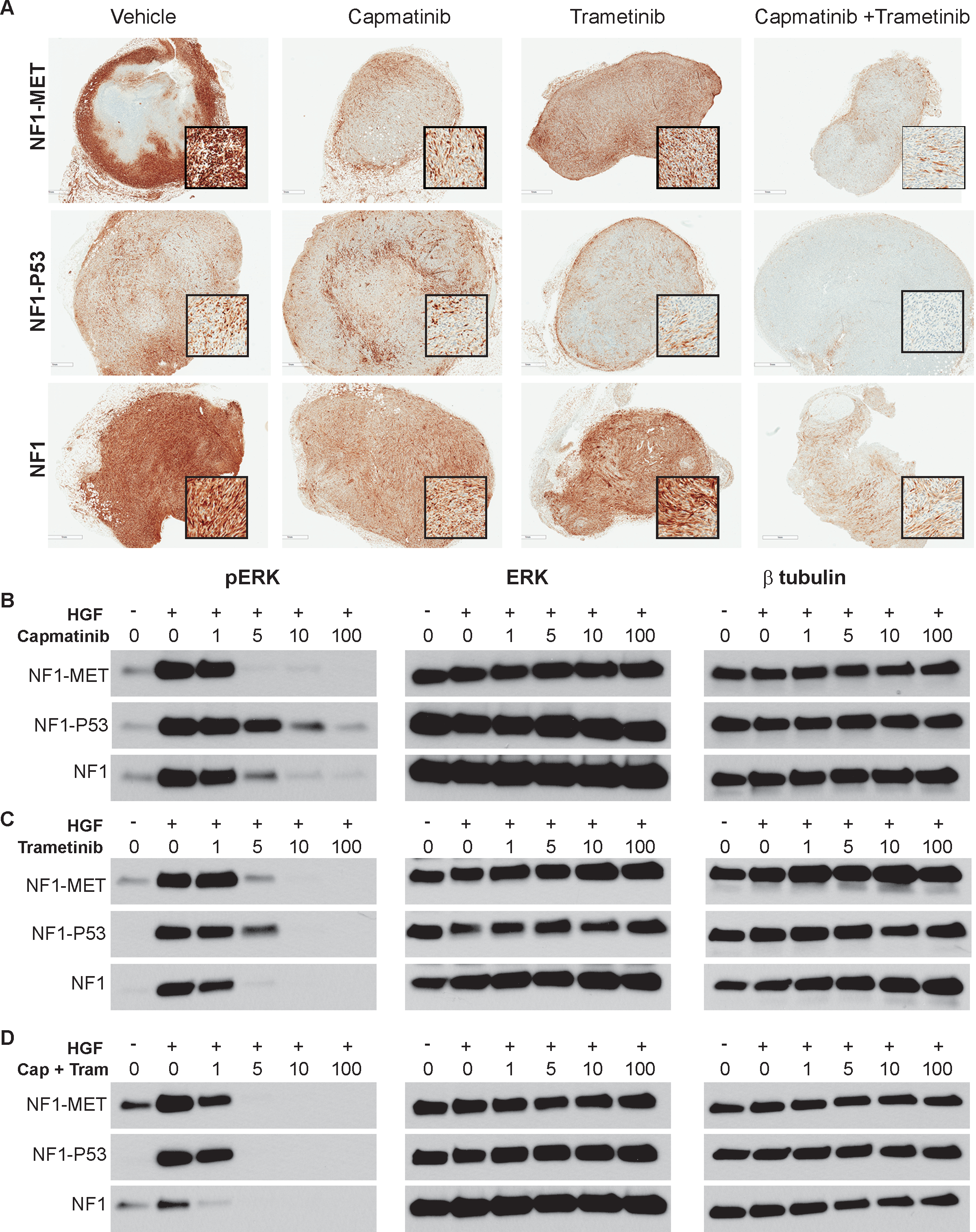
Responses in ERK signaling following MET and MEK inhibition in NF1 MPNST Models. (A) Immunostaining of pERK (T202/Y204) in MPNST models that were treated with kinase inhibitors for 2 days (areas of strongest signal are shown in the magnified inset boxes). (B-D) Western Blot analysis of MPNST cells treated with TKIs for 2 hours and +/-HGF for pERK (left panels), total ERK (middle panels), and β-tubulin control (right panels).

**Figure 6:**
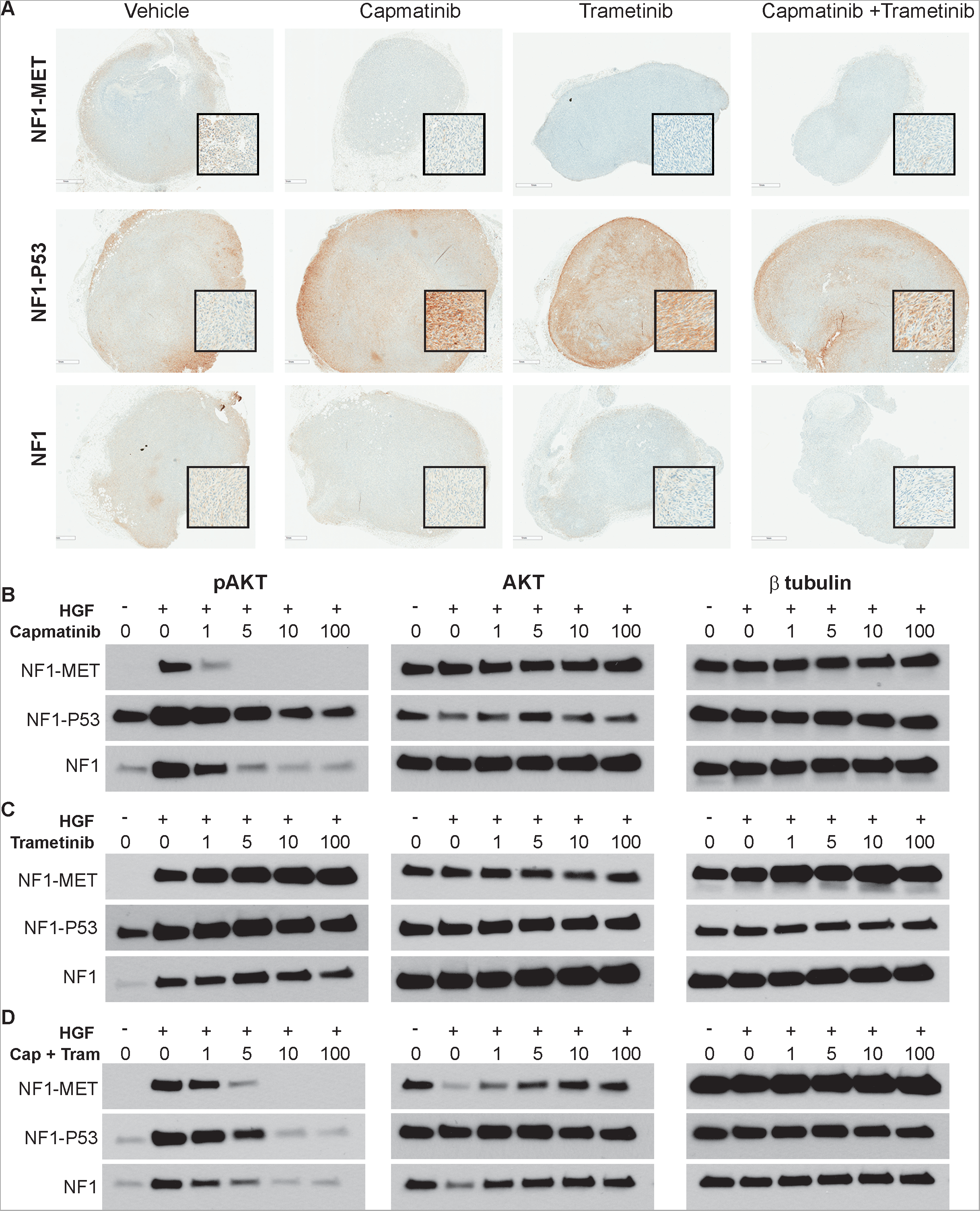
Responses in AKT signaling following MET and MEK inhibition of NF1-MET, NF1-P53, and NF1 MPNSTs. (A) Immunostaining of pAKT (S473) in MPNST models that were treated with kinase inhibitors for 2 days (areas of strongest signal are shown in the magnified inset boxes). (B-D) Western Blot analysis of MPNST cells treated with TKIs for 2 hours and +/-HGF for pAKT (left panels), total AKT (middle panels), and β-tubulin control (right panels).

### Differential response to MEK and MET inhibitors is associated with P53 status

We hypothesized that our MPNST models would demonstrate distinct growth patterns and kinome adaptation to MET and MEK inhibition. In NF1-MET (p53 wild type) MPNSTs we observed exquisite sensitivity to single agent capmatinib as well as significant tumor growth inhibition with trametinib (p < 1.0e^-16^; both capmatinib and trametinib treatments); however, pairwise comparison revealed that capmatinib was significantly more effective than trametinib in the NF1-MET tumors (p < 0.0004; Figure 1A-B). Examining the individual growth curves for each treatment vs. vehicle control groups allowed us to assess the variability in treatment response. Capmatinib treatment was associated with minimal response variability (SD=72 mm^3^) while trametinib treatment response variability (446 mm^3^) was substantially higher (Figure 1C). These treatment results suggest that NF1-MET tumors are “MET-addicted” and may have a distinctive kinome response to MET inhibition compared to other non-addicted models. We observed that capmatinib treatment resulted in complete inhibition of MET activation in NF1-MET tumors after 2 days of treatment (Figure 4A). Even though NF1-MET cells produce extremely high levels of MET compared to NF1-P53 and NF1 cells, 5 nM capmatinib inhibited pMET in the NF1-MET MPNST cells (Figure 4B). Interestingly, MET was strongly activated with trametinib treatment in the NF1-MET and NF1 tumorgrafts (Figure 4A) and in the NF1-MET, NF1-P53, and NF1 cells with HGF treatment (Figure 4C). In the NF1-MET tumors, trametinib treatment resulted in the expected ERK inactivation (Figure 5C); however AKT activation was drastically increased with trametinib treatment (Figure 6C). These results suggest that PI3K/AKT signaling may be a critical bypass mechanism in of MEK inhibition in MET-addicted MPNSTs. Moreover, MEK inhibition drives feedback activation of MET in NF1-related MPNSTs.

To understand how P53 loss affects tumor response to kinase inhibition, we used the NF1-P53 model (*NF1*^*KO/+*^*;p53*^*LSL-*R172H^), which recapitulates a p53^KO/+^ environment since the LSL cassette prevents expression of the *p53*^*R172H*^ mutant (15, 34). MPNST tumors from NF1-P53 mice have LOH of the wildtype P53 allele and we verified that the *p53*^*LSL-R172H*^ allele is in a *cis* conformation with *NF1* on Chr11 (15). *In vivo,* we observed that NF1-P53 tumors have a more aggressive growth rate (Figure 2A) than NF1-MET (Figure 1A) or NF1 tumors (Figure 3A). In NF1-P53 tumors capmatinib only partially inhibited tumor growth (Figure 2A-B; *p* <0.01) and there was substantial variability in response to capmatinib treatment (Figure 2C; SD=830 mm^3^).

Interestingly, there was no change in pMET after 2 days of capmatinib treatment in the NF1-P53 tumors despite the presence of *minor Met* amplification (Figure 4A). Trametinib had a substantial effect on NF1-P53 tumor growth (Figure 2A-B; *p* <0.007) yet significant response heterogeneity was observed (Figure 2C; SD=821 mm^3^). In pairwise comparison, both capmatinib and trametinib were significantly more effective as single agents than doxorubicin (Figure 2B) and reduced tumor growth by 35% and 39% respectively (Supplementary Figure 1B). Interestingly, there was no significant difference in response to single agent capmatinib vs. trametinib. Even though the response to single agent TKI was statistically significant, the NF1-P53 tumor growth curves still exhibited an aggressive growth pattern after 21 days of treatment. Another striking observation was that the NF1-MET and NF1 MPNST tumorgraft models demonstrated extremely high ERK activation compared to the moderate ERK activation observed in NF1-P53 tumors (Figure 5A, vehicle panel). Capmatinib substantially decreased pERK activation in both NF1-MET and NF1 tumors, but not in the NF1-P53 MPNSTs, whereas trametinib decreased ERK activity in all of the NF1 models (in comparison to vehicle, Figure 5A). In cells, trametinib inactivated ERK completely at 10 nM for NF1-MET and NF1-P53 cells, and at 5 nM in NF1 MPNST cells (Figure 5C). The variability in response to trametinib and the lower level of ERK activity in the NF1-P53 tumors suggests that these P53-deficient tumors may not be completely dependent on RAS-ERK signaling.

We also evaluated the efficacy of MET and MEK inhibition in P53 wild type, NF1 tumors which had a significantly slower growth rate compared to NF1-MET and NF1-P53 tumors (Figure 3A). NF1 tumor growth was suppressed with capmatinib (*p* < 0.0009) and trametinib (*p* < 1.0e^-16^), but was not significantly inhibited with doxorubicin (*p* < 0.07) (Figure 3A). In contrast to the MET-dependent NF1-MET tumor response, MEK inhibition with trametinib was more effective than MET inhibition in the NF1 tumors (Figure 3B, *p* < 0.0001). Interestingly, the RAS-MET response in NF1 tumors was distinct from the NF1-MET and NF1-P53 tumors. For example, capmatinib triggered AKT activation similar to NF1-MET tumors and ERK activation more similar to NF1-P35 tumors. Overall these results indicate that single agent therapy is less effective in P53-deficient (NF1-P53) tumors compared to P53 wild type tumors (NF1-MET, NF1).

### Combination MET and MEK inhibition is more efficacious than single agent regardless of P53 status

Due to the diverse bypass mechanisms of resistance to kinase inhibition, achieving a durable clinical response will likely require inhibition of multiple critical kinase signaling nodes. In NF1-MET tumors, combination therapy of capmatinib + trametinib and capmatinib + trametinib + doxorubicin resulted in an overall growth rate reduction of 60.3% and 67.0% respectively (Supplementary Figure 1A). Even though combination therapy of capmatinib + trametinib (65 mm^3^) or capmatinib + trametinib + doxorubicin (19 mm^3^) did minimize the variability in response compared to capmatinib alone (72 mm^3^) (Figure 1C), there was no significant improvement in tumor reduction with combination therapy in the NF1-MET tumors as shown by pair-wise comparison (Figure 1B). The reduction in growth rates relative to vehicle showed a 75.7% reduction with capmatinib alone, 94.5% with capmatinib + trametinib, and a 103.8% growth rate reduction with capmatinib + trametinib + doxorubicin (Supplementary Figure 1A). These results indicate that NF1-related MPNSTs containing *MET* amplification are highly sensitive to MET inhibitors, yet clinical response may be further improved with the addition of a MEK inhibitor.

In NF1-P53 tumors, combined capmatinib + trametinib treatment significantly inhibited tumor growth (p < 0.000007; Figure 2A), yet the addition of doxorubicin did not significantly improve tumor inhibition (cap+tram+dox vs. cap+tram, p < 0.7). Analysis of the individual growth curves for NF1-P53 tumors revealed drastic improvement in response variability with combination therapy compared to single agent alone (Figure 2C). Capmatinib + trametinib reduced the variability in response to 638 mm^3^, yet there was one ‘non-responder’ that impacted this variability, whereas the triple combination of capmatinib + trametinib + doxorubicin had the smallest variability (344 mm^3^). These results confirm that NF1-related MPNSTs with P53 deficiency are less responsive to therapeutic approaches targeting single kinases and the best therapeutic response is achieved by inhibiting both MET and MEK signaling.

In the NF1 tumors, capmatinib + trametinib showed statistically significant improvement in tumor suppression vs single agent therapy with either trametinib or capmatinib, but no benefit was observed with the addition of doxorubicin to the capmatinib + trametinib combination (Figure 3B, *p* < 0.07). Overall, there was less heterogeneity in treatment response in the NF1 tumors (Figure 3C) compared to the NF1-MET (Figure 1C) and NF1-P53 (Figure 2C) tumors. Combination treatment with capmatinib + trametinib resulted in a very uniform response (SD= 39 mm^3^) compared to single agent treatment of capmatinib (SD=159 mm^3^) and trametinib (130 mm^3^) (Figure 3C). The reduction in tumor growth rate was similar between capmatinib + trametinib (110%) and the capmatinib + trametinib + doxorubicin (108%) treatments (Supplementary Figure 1C). These results indicate that NF1-related MPNSTs without *Met* amplification or *p53* loss may also be responsive to combination MET and MEK inhibition.

### Combination MET and MEK inhibition prevents adaptive ERK and AKT response

Differential patterns of ERK inactivation and activation were observed in response to monotherapy vs. combination kinase inhibition in the NF1 models. Overall, combined MET and MEK inhibition resulted in a greater than 90% reduction in MET and ERK activation in all of the MPNST models (Figure 4A and Figure 5A, right panels). In NF1-MET cells, ERK activation was significantly inhibited by 5 nM capmatinib, whereas NF1 and especially NF1-P53 cells showed a marginal decrease in pERK in response to capmatinib (Figure 5B). Trametinib was much more effective at inhibiting ERK activation and demonstrated reduction of pERK at 10nM in NF1-P53 and NF1-MET cells and at 5nM in NF1 cells (Figure 5C). Combined capmatinib and trametinib treatment mirrored the tumorgraft results (Figure 5A) and was the most effective treatment in decreasing pERK (Figure 5D). These findings validate the expected inhibition of RAS/ERK signaling with MEK TKIs. Moreover, these results emphasize the importance of combined kinase inhibition to eradicate RAS signaling (Figures 1-3).

Another major signaling pathway that is activated by MET and known to interact with RAF is the PI3K/AKT pathway. We observed high pAKT at the invasive edges of the NF1-P53 tumors, yet minimal pAKT was present in NF1-MET or NF1 tumors (Figure 6A). Interestingly, increased pAKT was observed in NF1-P53 tumors after capmatinib or trametinib treatment and persisted at the tumor periphery even with combined capmatinib and trametinib treatment (Figure 6A). Analysis of AKT activity in the MPNST lines verified our *in vivo* observations. Even though AKT expression is comparable across the MPNST lines, pAKT is significantly higher in the NF1-P53 cells at basal conditions (Figures 6B-D). As expected, MET activation with HGF treatment stimulates pAKT in NF1-MET and NF1 cells, and further augmented AKT activation in NF1-P53 cells. Capmatinib treatment resulted in strong inhibition of AKT in NF1-MET and NF1 cells, but was ineffective in diminishing pAKT in the NF1-P53 cells (Figure 6B). Trametinib treatment resulted in a significant increase in pAKT in all three of the MPNST cells (Figure 6C). Combined MET and MEK inhibition was the most effective treatment for abrogating AKT activation in the NF1-P53 cells (Figure 6D). This analysis reveals that NF1-P53 tumors have basal AKT activity that is rapidly increased upon MET or MEK inhibition. This may indicate a distinctive mechanism of resistance that is readily activated upon kinase inhibition in P53-deficient MPNSTs.

### MEK inhibitor resistance is mediated through RAS-MET signaling leading to sustained AKT activation

We observed that trametinib treatment activated MET in a dose-, HGF-dependent manner in NF1-MET cells (Figure 4C). The *Met* amplified, NF1-P53 and *Hgf*-amplified NF1 lines also demonstrated sustained MET signaling within the therapeutic trametinib range. The activation of MET following trametinib treatment was unanticipated, but suggests that feedback RTK activation occurs following RAS blockade in NF1-deficient cells. Even though NF1-MET cells expressed substantially more MET than the NF1-P53 and NF1 cells at baseline (Figure 4B), HGF-induced MET activation was eliminated with 5 nM in the NF1-MET cells. As observed in the tumorgrafts (Figure 4A), dual MET and MEK inhibition effectively abrogated pMET levels in all three lines at 1 nM capmatinib (Figure 4D) and resulted in a significant reduction in tumor growth compared to single agent alone (Figures 1-3). As mentioned earlier, trametinib treatment resulted in increased pAKT in all three of the MPNST cells. Although combined MET and MEK inhibition suppressed AKT activation to its non-stimulated state, pAKT was not completely inhibited in the NF1-P53 and NF1 lines (Figure 6D). These observations strongly point to a robust compensation mechanism that allows for sustained AKT signaling despite abrogation of MET and MEK signaling. These results also confirm that MEK inhibition promotes MET activation, particularly when genomic HGF or MET alterations are present.

### RAF reinforces RAS-AKT crosstalk in MPNSTs

An alternative explanation for trametinib-based activation of MET or sustained AKT activation with combination MET-MEK inhibition is that the drugs themselves modulate RAS-MET signaling independent of HGF. In order to address this question, and to assess the impact of capmatinib and trametinib therapy on the broader kinome, we performed experiments where we varied single agent and combination therapy concentrations, with and without HGF stimulation. We examined both upstream regulatory proteins (BRAF/CRAF, MET) and downstream RAS-MET effector phosphoproteins (AKT, STAT3, ERK). The impact of P53 status was also analyzed by using the P53-intact, NF1-MET lines and P53-deficient NF1-P53 lines. AKT was activated at baseline in the NF1-P53 lines, but not NF1-MET (Figure 7A-B). HGF strongly induced AKT phosphorylation in both NF1-MET and NF1-P53 cells, an effect that was inhibited by capmatinib but not fully abrogated in the context of P53 deficiency (Figure 7B). MEK inhibition with trametinib did not activate AKT independent of HGF in the NF1-MET line, but strong activation above the HGF-stimulated state was noted when trametinib and HGF were combined (Figure 7A). In the NF1-P53 cells, trametinib activated AKT above baseline without HGF and strongly activated AKT with HGF above the stimulated state. Combined capmatinib + trametinib treatment reduced pAKT levels, yet pAKT was increased compared to baseline with the combination therapy (Figure 7B). These results confirm that both HGF-dependent and non HGF-dependent activation of AKT occurs through MET with trametinib treatment.

**Figure 7:**
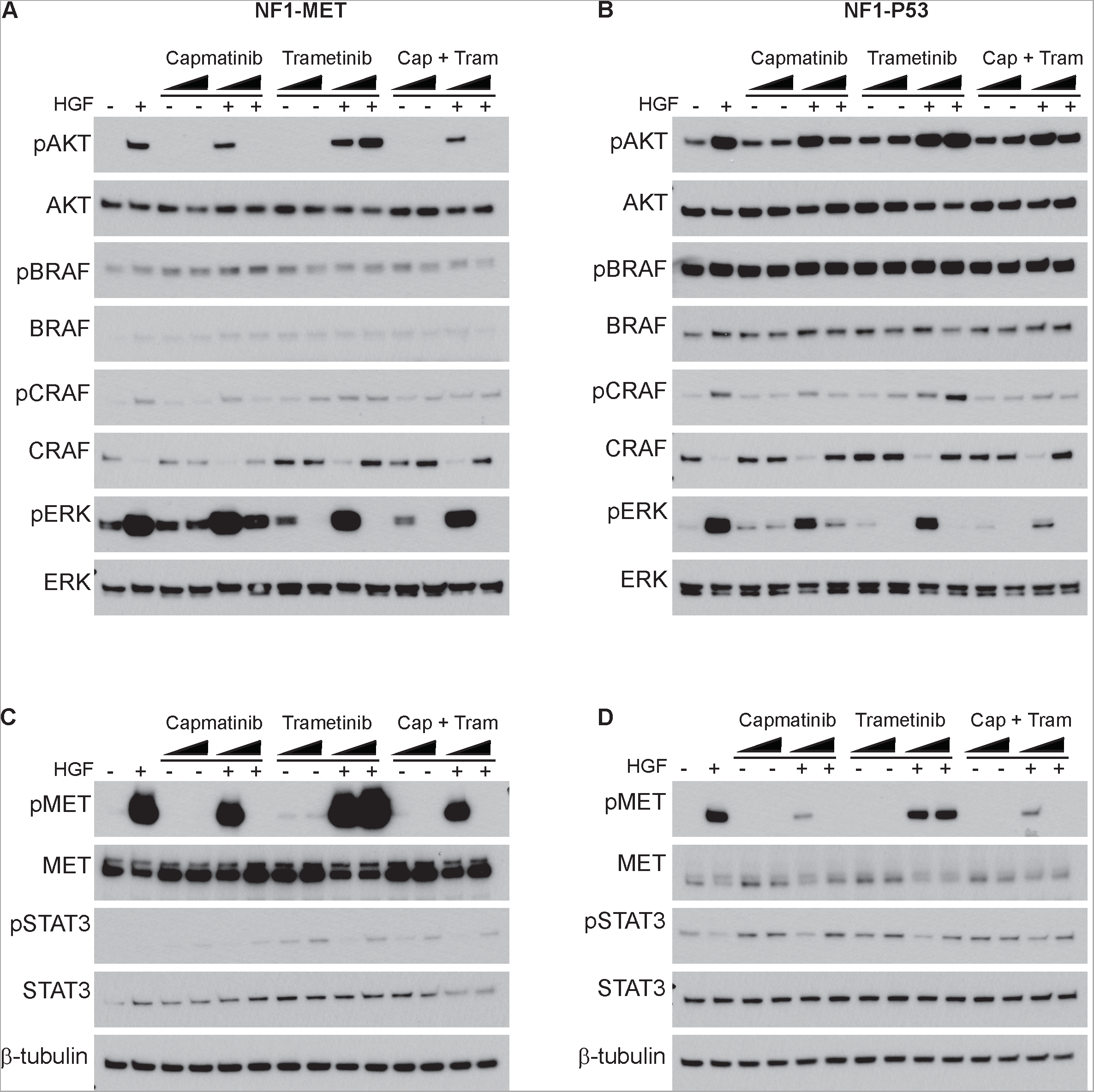
RAF reinforces RAS-AKT crosstalk in NF1-MET and NF1-P53 MPNSTs. (A-B) Western Blot analysis of NF1-MET (A) and NF1-P53 (B) MPNST cells treated with TKIs for 2 hours and +/-HGF for AKT (S473), RAF (Ser445), and ERK (T202/Y204) activation. At the low and high dose ranges (represented by dark triangle), cells were treated with 1 nM or 100 nM of capmatinib and/or trametinib respectively. (C-D) Western blot analysis of NF1-MET (C) and NF1-P53 (D) MPNST cells for MET (Y1234/1235) and STAT3 (Y705). β-tubulin was run as a control for all lysates.

One of the targetable mediators between the RAS and PI3K/AKT pathways is RAF (35, 36). Low BRAF expression was observed in NF1-MET cells compared to NF1-P53 cells and BRAF activation was not affected by MET or MEK inhibition (Figure 7A-B). Capmatinib was associated with mild activation of bRAF above basal state in the NF1-MET lines; however no change was noted in the NF1-P53 lines possibly due to saturation from overexpression.

Increase in pCRAF was observed with HGF treatment in both the NF1-MET and NF1-P53 cells. Interestingly, capmatinib inhibited the effects of HGF stimulation on cRAF phosphorylation, but trametinib increased p-CRAF (in the presence of HGF), an effect that was mitigated by combined MET and MEK inhibition (Figure 7A-B). We observed a correlative pattern of total CRAF expression and activation similar to the relationship that is observed with RTK activation and receptor turnover. We also evaluated STAT signaling and observed a decrease in STAT3 activation in the presence of HGF in NF1-P53 cells but no change in NF1-MET cells (Figure 7C-D). However, in NF1-P53 cells, pSTAT3 increased in response to monotherapy of both capmatinib and trametinib and also combined capmatinib+trametinib treatment (Figure 7D). This was also observed to a weaker extent in NF1-MET cells (Figure 7C). Overall, this analysis of the RAS-AKT kinome reveals that P53 deficiency is associated with a high degree of baseline AKT activation that serves as a compensation point to promote survival in response to combined MET and MEK inhibition. Moreover, the data also supports the concept that 1) AKT activation is regulated through RAF and 2) AKT is activated by MEK inhibition but not MET inhibition. These findings identify key nodes that drive kinome robustness in the context of MET-addiction and P53 deficiency and reveal the inherent signaling feedback mechanisms that promote drug resistance in MPNSTs.

## Discussion

RAS-MET signal interactions are emerging as important mechanisms of therapy resistance and disease progression. NF1-related MPNSTs are no exception as they exhibit both MET and *HGF* amplifications, and deregulated RAS signaling owing to loss of NF1-mediated tumor suppression. To date, the therapeutic implication of these cytogenetic alterations has not been established. Based on the data presented above, highly selective and potent MET and MEK inhibitors are available for clinical use and are promising therapies for MPNSTs. However, in order to effectively use tyrosine kinase inhibitors in MPNST treatment it will be critical to not only inhibit oncogenic RTK signaling, but also anticipate the adaptive kinome response that occurs through the RAS or mTOR pathways. The mitigated therapy response observed in other RAS-deregulated cancers following use of single agent kinase inhibitors (i.e. EGFR inhibition in non– small cell lung cancer) teaches us valuable lessons about kinome adaptation. As in KRAS mutant lung cancer, it is clear that NF1-deficient MPNSTs also rely on functionally redundant kinome signaling networks, not just the driver effects of a single therapeutic target (30, 31, 37). Combination therapies, therefore, are better suited to target the diversity of kinase signaling pathways that drive disease progression. In order to address this concept in MPNSTs, we 1) determined the efficacy and kinome response to MET and/or MEK inhibition, 2) evaluated how P53 modulates kinome signaling, and 3) examined targetable points of signaling convergence between the RAS and MET pathways. Taken together, our data supports the concept that RAS-MET signal interactions are both essential to disease progression and targetable.

We tested the efficacy of single agent and combined inhibition of MET and MEK in MPNST models with variable levels of *MET* amplification and *P53* deficiency. We discovered that NF1-MET tumors are “MET-addicted”, extremely sensitive to capmatinib treatment, and have a unique kinome response to MET inhibition in comparison to non-*MET* amplified MPNSTs. Even though NF1-MET tumors were highly sensitive to MET inhibition, response heterogeneity was decreased with combined MET and MEK inhibition. Trametinib was the most effective signal agent therapy in both NF1-P53 and NF1 tumors, yet combined capmatinib + trametinib treatment was significantly more effective than single agent therapy in these NF1-P53 and NF1 tumors. Interestingly, doxorubicin was only marginally effective in NF1 tumors and resulted in a significantly worse response compared in capmatinib or trametinib in all of the MPNST models. Overall, our results demonstrated that NF1-related MPNSTs have the best response to combined MET and MEK inhibition regardless of the status of *MET* and *P53*.

By evaluating the MET and RAS signaling response to MET and/or MEK inhibition in our MPNST models, we were able to identify distinct kinome responses to MET or MEK inhibition. We observed a differential pattern of kinase signaling between the P53-intact and P53-deficient models, and in response to MET and MEK inhibition. We also observed consistent patterns of kinome adaptation to single agent MET or MEK inhibition that could be applicable to other disease contexts as they occurred agnostic to P53 status, and ultimately led to therapy resistance in our models. In all of the MPNST models, combined MET and MEK inhibition was the most effective treatment for decreasing ERK activity. The fact that NF1-P53 tumors have a lower basal ERK activity and a variable response to MEK inhibition suggests that P53-deficient tumors may not be exclusively dependent on RAS-ERK signaling. This idea is supported by high basal AKT activity in NF1-P53 tumors that was rapidly increased upon MET or MEK inhibition. Given that we observed increase AKT activation in response to trametinib in all of our MPNST tumor models, PI3K/AKT pathway may be a robust compensation mechanism in RAS-deregulated MPNSTs. As with ERK, combined MET and MEK inhibition was the most effective treatment for decreasing activation of AKT. We did not evaluate all of the cross-talk candidates that could potentially activate AKT in the setting of RAS-MET inhibition, but our data suggest that both the BRAF and CRAF may play a role in activating AKT in response to MET or MEK inhibition.

Unexpectedly, we observed that trametinib treatment resulted in MET activation in all of the NF1 MPNST models. Trametinib-mediated MET activation was dependent on HGF and could be abrogated with the addition of capmatinib. Taken together, these data confirm that distinct compensation mechanisms drive resistance to capmatinib and trametinib therapy, and broadly implicate HGF/MET signal activation in resistance to MEK inhibition. Even though RAS signaling is not exclusively activated by MET, MET expression is significantly upregulated in RAS-transformed cells and is required for RAS-mediated tumorigenesis and metastasis (38, 39). Observed resistance to mTOR inhibitors in breast, lung, and renal cell cancer is attributable to MET activation (40), an effect that can be overcome through addition of a MET inhibitor (41). These data strongly point towards signaling convergence between the RAS and MET pathways in cancer and highlight a potential bi-directional mechanism of RAS-MET activation in the setting of tyrosine kinase inhibition and ligand-dependent activation of RAS.

There is an evolving conceptual framework that implicates P53 as a key regulatory protein for kinase signaling, either as an indirect driver of adaptive genomic alterations such as RTK amplifications, or through transcriptional regulation of oncogenic proteins such as VEGFR or heat-shock factor 1 (HSF1) leading to EGFR and ErbB2 signal activation (42, 43). P53 haploinsufficiency was recently shown to cooperate with *EGFR* amplification to increase tumor grade in murine MPNSTs (44). Another example of the connection between P53 and kinase signaling is the recent observation that mammary stem cell expansion in triple-negative breast cancer was shown to be dependent on P53 deficiency (23). In this same study, MET activation was a direct consequence of P53 deficiency and promoted robust maintenance of the stem cell phenotype. Our data supports the integral role that P53 status in manipulating the kinome in cancer progression and therapeutic resistance.

In conclusion, this work extends our initial findings that MET is sufficient for malignant transformation of NF1-deficient peripheral nerve cells into MPNSTs and can be successfully targeted with a highly specific MET inhibitor (15). Our demonstration of continued MET dependency following transformation and in the setting of MEK inhibition demonstrates the kinome adaptations that may exist in human MPNSTs. Our work verifies the significant role that MET signaling plays in reinforcing states of RAS activation and the ability of the RAS/ERK pathway to reinforce MET signaling. Based on the data presented, it is possible that MEK targeted therapy strategies can inadvertently activate MET resulting in broader TKI resistance. MET compensation should be assessed when analyzing the therapeutic response to TKI’s in RAS-addicted tumors.

## Methods

### Histopathology

Mouse tissues were fixed in 10% neutral buffered formalin for 72 hours (Thermo) prior to paraffin embedding and sectioning for histology and immunohistochemistry.

Immunohistochemistry was performed on formalin-fixed paraffin embedded samples using a citrate-based antigen retrieval system (Vector Labs). Samples were stained for phosphorylated MET (Cell Signaling D26 #3077), phosphorylated MAPK (Cell Signaling #9101), phosphorylated AKT (Cell signaling D9E #4060), and phosphorylated MEK (Cell Signaling 166F8 #2338) were performed using a Ventana autostainer. Images were obtained with an Aperio Digital Imaging system (Leica).

### Western Blotting

Mouse derived cell lines NF1-p53, NF1-MET, and NF1 were grown to 90% confluency overnight by seeding plates with 500,000 – 750,000 cells. Cells were then serum starved overnight and then treated with indicated inhibitors (capmatinib, trametinib, or combination) for 2 hours followed by 100ng/mL of HGF for 15 minutes. Cells were then washed with PBS and harvested in RIPA buffer plus protease inhibitor cocktail (Roche). Lysates (16-20 μg) were resolved on a 4%-20% TGX SDS-PAGE gel (Bio-Rad) and transferred to a PVDF membrane (Invitrogen). After blocking for 1 hour with 5% dry milk in TBST buffer (20 mmol/L Tris-HCl pH 7.4, 150 mmol/L NaCl, 0.1% Tween-20), blots were probed overnight at 4°C with the following primary antibodies from Cell Signaling Technology: Met (25H2; #3127), pMET (D26; #3077), AKT (#9272), pAKT (S473; #9271), MAPK (#9102), pMAPK (Thr202/Tyr204; #9101), p-CRAF (#9427), p-BRAF(#2696), CRAF (#12552), BRAF (#9433), pSTAT3 (#9145), STAT3 (#9139), and β-tubulin (#2146). Blots were reacted with peroxidase-conjugated secondary antibody for 1 hour (Cell Signaling Technology) and protein bands were visualized using ECL detection (Amersham).

### Development of murine MPNST tumorgrafts and *in vivo* testing of TKIs

Immediately following euthanasia of tumor bearing mice, 15-25 mg portions of each tumor were transplanted into the flank of four gender-matched athymic nude recipient mice using a 10 gauge trochar (Innovative Research of America, Sarasota FL). Mice were examined weekly and euthanized when the tumor size exceeded 1500mm3 by external caliper measurement.

These tumors were then passed into new immunocompromised mice for treatment studies. Bulk tumor pieces were transplanted subcutaneously into athymic nude female mice and tumor growth was evaluated twice weekly. When tumor volume reached approximately 150 mm^3^, mice were randomized into treatment groups and dosed 5 days per week for 3 weeks or until mice reached euthanasia criteria for tumor burden (tumor volume >2500mm^3^). Respective doses across all treatment combinations were Capmatinib 30mg/kg twice daily via oral gavage, Trametinib 1 mg/kg daily via oral gavage, and Doxorubicin 1 mg/kg twice weekly via intraperitoneal injection. Capmatinib and Trametinib were obtained from Novartis. The tumors were measured twice weekly using a caliper and the tumor volumes were calculated as length x width x depth.

### Statistical Analysis

Linear mixed-effects models with false discovery rate adjusted contrasts were used to estimate and compare tumor growth rates between different treatment combinations within each tumor line. 10,000 non-parametric bootstrap resampled datasets were used to estimate the confidence intervals for the percent reduction in tumor growth rates. All analyses were conducted using Rv3.2.2 (https://cran.r-project.org/) with an assumed level of significance of α = 0.05.

## Authors’ Contributions

**Conception and design:** CR Graveel, MR Steensma

**Development of methodology:** MG Pridgeon, CJ Essenburg

**Acquisition of data:** MG Pridgeon, EA Tovar, CJ Essenburg, PS Dischinger

**Analysis and interpretation of data:** Z Madaj, MG Pridgeon, EA Tovar, CR Graveel, MR Steensma

**Writing, review, and/or revision of the manuscript:** MG Pridgeon, CR Graveel, MR Steensma

**Administrative, technical, or material support:** JD Peacock, F Maina, R Dono

**Study supervision:** MR Steensma

## Acknowledgements

We would like to thank the Lisa Turner and the VARI Histology Core for their expertise and the VARI Vivarium for their continuous dedication.

## Supplemental Figure Legends

**Figure S1:**
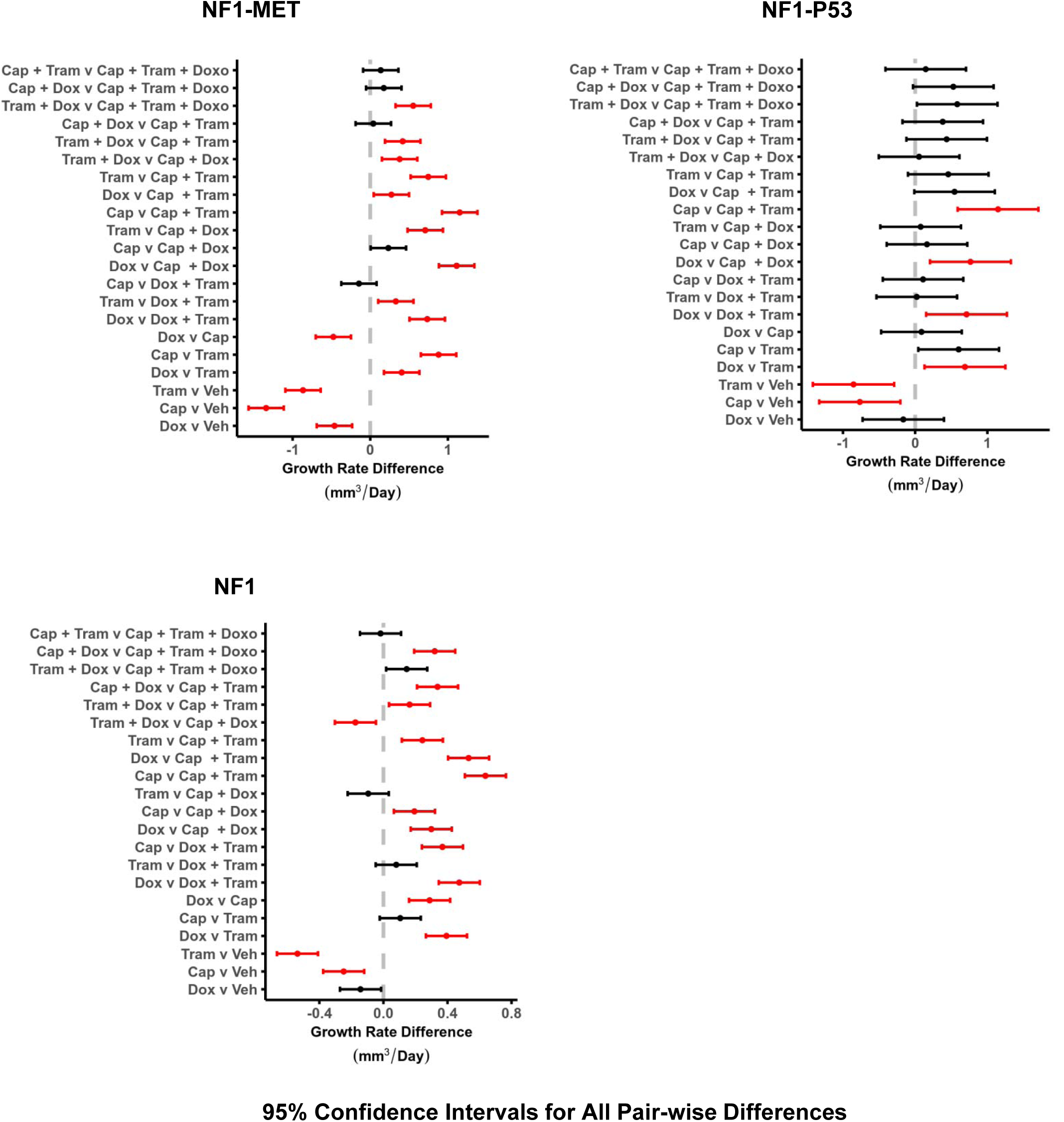
Mean percent reduction in growth rates of mice treated with each treatment arm versus vehicle. Error bars are the bootstrap 95th percentile interval, estimated based on 10,000 resampled datasets.

## References

1. Lammert, M., Friedman, J.M., Kluwe, L., and Mautner, V.F. 2005. Prevalence of neurofibromatosis 1 in German children at elementary school enrollment. Arch Dermatol 141:71–74.

2. Evans, D.G., Howard, E., Giblin, C., Clancy, T., Spencer, H., Huson, S.M., and Lalloo, F. 2010. Birth incidence and prevalence of tumor-prone syndromes: estimates from a UK family genetic register service. Am J Med Genet A 152A:327-332.

3. Wallace, M.R., Marchuk, D.A., Andersen, L.B., Letcher, R., Odeh, H.M., Saulino, A.M., Fountain, J.W., Brereton, A., Nicholson, J., Mitchell, A.L., et al. 1990. Type 1 neurofibromatosis gene: identification of a large transcript disrupted in three NF1 patients. Science 249:181–186.

4. Friedrich, R.E., Hartmann, M., and Mautner, V.F. 2007. Malignant peripheral nerve sheath tumors (MPNST) in NF1-affected children. Anticancer Res 27:1957–1960.

5. Ingham, S., Huson, S.M., Moran, A., Wylie, J., Leahy, M., and Evans, D.G. 2011. Malignant peripheral nerve sheath tumours in NF1: improved survival in women and in recent years. Eur J Cancer 47:2723–2728.

6. Evans, D.G., Baser, M.E., McGaughran, J., Sharif, S., Howard, E., and Moran, A. 2002. Malignant peripheral nerve sheath tumours in neurofibromatosis 1. J Med Genet 39:311–314.

7. Porter, D.E., Prasad, V., Foster, L., Dall, G.F., Birch, R., and Grimer, R.J. 2009. Survival in Malignant Peripheral Nerve Sheath Tumours: A Comparison between Sporadic and Neurofibromatosis Type 1-Associated Tumours. Sarcoma 2009:756395.

8. Watson, K.L., Al Sannaa, G.A., Kivlin, C.M., Ingram, D.R., Landers, S.M., Roland, C.L., Cormier, J.N., Hunt, K.K., Feig, B.W., Ashleigh Guadagnolo, B., et al. 2017. Patterns of recurrence and survival in sporadic, neurofibromatosis Type 1-associated, and radiation-associated malignant peripheral nerve sheath tumors. J Neurosurg 126:319–329.

9. Fukuda, T., Ichimura, E., Shinozaki, T., Sano, T., Kashiwabara, K., Oyama, T., Nakajima, T., and Nakamura, T. 1998. Coexpression of HGF and c-Met/HGF receptor in human bone and soft tissue tumors. Pathol Int 48:757–762.

10. Mantripragada, K.K., Spurlock, G., Kluwe, L., Chuzhanova, N., Ferner, R.E., Frayling, I.M., Dumanski, J.P., Guha, A., Mautner, V., and Upadhyaya, M. 2008. High-Resolution DNA Copy Number Profiling of Malignant Peripheral Nerve Sheath Tumors Using Targeted Microarray-Based Comparative Genomic Hybridization. Clinical Cancer Research 14:1015–1024.

11. Torres, K.E., Zhu, Q.-S., Bill, K., Lopez, G., Ghadimi, M.P., Xie, X., Young, E.D., Liu, J., Nguyen, T., Bolshakov, S., et al. 2011. Activated MET Is a Molecular Prognosticator and Potential Therapeutic Target for Malignant Peripheral Nerve Sheath Tumors. Clinical Cancer Research 17:3943–3955.

12. Gherardi, E., Birchmeier, W., Birchmeier, C., and Vande Woude, G. 2012. Targeting MET in cancer: rationale and progress. Nat Rev Cancer 12:89–103.

13. Straussman, R., Morikawa, T., Shee, K., Barzily-Rokni, M., Qian, Z.R., Du, J., Davis, A., Mongare, M.M., Gould, J., Frederick, D.T., et al. 2012. Tumour micro-environment elicits innate resistance to RAF inhibitors through HGF secretion. Nature 487:500–504.

14. Oddo, D., Siravegna, G., Gloghini, A., Vernieri, C., Mussolin, B., Morano, F., Crisafulli, G., Berenato, R., Corti, G., Volpi, C.C., et al. 2017. Emergence of MET hyper-amplification at progression to MET and BRAF inhibition in colorectal cancer. Br J Cancer 117:347–352.

15. Peacock, J.D., Pridgeon, M.G., Tovar, E.A., Essenburg, C.J., Bowman, M., Madaj, Z., Koeman, J., Grit, J.L., Dodd, R.D., Cardona, D.M., et al. 2017. Genomic MET amplification occurs early in NF1-related malignant peripheral nerve sheath tumor (MPNST) progression and is a potent therapeutic target. submitted.

16. Upadhyaya, M., Spurlock, G., Majounie, E., Griffiths, S., Forrester, N., Baser, M., Huson, S.M., Gareth Evans, D., and Ferner, R. 2006. The heterogeneous nature of germline mutations in NF1 patients with malignant peripheral serve sheath tumours (MPNSTs). Human Mutation 27:716–716.

17. Miller, S.J., Rangwala, F., Williams, J., Ackerman, P., Kong, S., Jegga, A.G., Kaiser, S., Aronow, B.J., Frahm, S., Kluwe, L., et al. 2006. Large-Scale Molecular Comparison of Human Schwann Cells to Malignant Peripheral Nerve Sheath Tumor Cell Lines and Tissues. Cancer Research 66:2584–2591.

18. Cichowski, K., Shih, T., Schmitt, E., Santiago, S., Reilly, K., McLaughlin, M., Bronson, R., and Jacks, T. 1999. Mouse models of tumor development in neurofibromatosis type 1. Science 286:2172–2176.

19. Kourea, H., Cordon-Cardo, C., Dudas, M., Leung, D., and Woodruff, J. 1999. Expression of p27(kip) and other cell cycle regulators in malignant peripheral nerve sheath tumors and neurofibromas: the emerging role of p27(kip) in malignant transformation of neurofibromas. Am J Pathol 155:1885–1891.

20. Lee, W., Teckie, S., Wiesner, T., Ran, L., Prieto Granada, C.N., Lin, M., Zhu, S., Cao, Z., Liang, Y., Sboner, A., et al. 2014. PRC2 is recurrently inactivated through EED or SUZ12 loss in malignant peripheral nerve sheath tumors. Nat Genet 46:1227–1232.

21. Buchstaller, J., McKeever, P.E., and Morrison, S.J. 2012. Tumorigenic cells are common in mouse MPNSTs but their frequency depends upon tumor genotype and assay conditions. Cancer Cell 21:240–252.

22. Tao, L., Xiang, D., Xie, Y., Bronson, R.T., and Li, Z. 2017. Induced p53 loss in mouse luminal cells causes clonal expansion and development of mammary tumours. Nat Commun 8:14431.

23. Chiche, A., Moumen, M., Romagnoli, M., Petit, V., Lasla, H., Jezequel, P., de la Grange, P., Jonkers, J., Deugnier, M.A., Glukhova, M.A., et al. 2017. p53 deficiency induces cancer stem cell pool expansion in a mouse model of triple-negative breast tumors. Oncogene 36:2355–2365.

24. Liu, W.T., Jing, Y.Y., Yu, G.F., Chen, H., Han, Z.P., Yu, D.D., Fan, Q.M., Ye, F., Li, R., Gao, L., et al. 2016. Hepatic stellate cell promoted hepatoma cell invasion via the HGF/c-Met signaling pathway regulated by p53. Cell Cycle 15:886–894.

25. Yang, X.P., Liu, S.L., Xu, J.F., Cao, S.G., Li, Y., and Zhou, Y.B. 2017. Pancreatic stellate cells increase pancreatic cancer cells invasion through the hepatocyte growth factor /c-Met/survivin regulated by P53/P21. Exp Cell Res 357:79–87.

26. Ma, B., Bang, Y.J., Lim, W.T., Nam, D.H., Su, W.C., Schellens, J.H.M., Azaro, A., Akimov, M., Zhang, Y., Kumar, A., et al. 2015. P1.04Phase I dose escalation and expansion study to evaluate safety and efficacy of INC280 in patients with advanced MET-dependent solid tumors. Annals of Oncology 26:ii17–ii17.

27. Bang, Y.-J., Su, W.-C., Nam, D.-H., Lim, W.-T., Bauer, T.M., Brana, I., Poon, R.T.-P., Hong, D.S., Lin, C.-C., Peng, B., et al. 2014. Phase I study of the safety and efficacy of INC280 in patients with advanced MET-dependent solid tumors. Journal of Clinical Oncology 32:2520–2520.

28. Liu, X., Wang, Q., Yang, G., Marando, C., Koblish, H.K., Hall, L.M., Fridman, J.S., Behshad, E., Wynn, R., Li, Y., et al. 2011. A novel kinase inhibitor, INCB28060, blocks c-MET-dependent signaling, neoplastic activities, and cross-talk with EGFR and HER-3. Clin Cancer Res 17:7127–7138.

29. Linklater, E.S., Tovar, E.A., Essenburg, C.J., Turner, L., Madaj, Z., Winn, M.E., Melnik, M.K., Korkaya, H., Maroun, C.R., Christensen, J.G., et al. 2016. Targeting MET and EGFR crosstalk signaling in triple-negative breast cancers. Oncotarget.

30. Wilson, T.R., Fridlyand, J., Yan, Y., Penuel, E., Burton, L., Chan, E., Peng, J., Lin, E., Wang, Y., Sosman, J., et al. 2012. Widespread potential for growth-factor-driven resistance to anticancer kinase inhibitors. Nature 487:505–509.

31. Chong, C.R., and Janne, P.A. 2013. The quest to overcome resistance to EGFR-targeted therapies in cancer. Nat Med 19:1389–1400.

32. Dombi, E., Baldwin, A., Marcus, L.J., Fisher, M.J., Weiss, B., Kim, A., Whitcomb, P., Martin, S., Aschbacher-Smith, L.E., Rizvi, T.A., et al. 2016. Activity of Selumetinib in Neurofibromatosis Type 1-Related Plexiform Neurofibromas. N Engl J Med 375:2550–2560.

33. Gilmartin, A.G., Bleam, M.R., Groy, A., Moss, K.G., Minthorn, E.A., Kulkarni, S.G., Rominger, C.M., Erskine, S., Fisher, K.E., Yang, J., et al. 2011. GSK1120212 (JTP-74057) is an inhibitor of MEK activity and activation with favorable pharmacokinetic properties for sustained in vivo pathway inhibition. Clin Cancer Res 17:989–1000.

34. Olive, K.P., Tuveson, D.A., Ruhe, Z.C., Yin, B., Willis, N.A., Bronson, R.T., Crowley, D., and Jacks, T. 2004. Mutant p53 gain of function in two mouse models of Li-Fraumeni syndrome. Cell 119:847–860.

35. Kim, J.Y., Welsh, E.A., Fang, B., Bai, Y., Kinose, F., Eschrich, S.A., Koomen, J.M., and Haura, E.B. 2016. Phosphoproteomics Reveals MAPK Inhibitors Enhance MET- and EGFR-Driven AKT Signaling in KRAS-Mutant Lung Cancer. Mol Cancer Res 14:1019–1029.

36. Byeon, H.K., Na, H.J., Yang, Y.J., Kwon, H.J., Chang, J.W., Ban, M.J., Kim, W.S., Shin, D.Y., Lee, E.J., Koh, Y.W., et al. 2016. c-Met-mediated reactivation of PI3K/AKT signaling contributes to insensitivity of BRAF(V600E) mutant thyroid cancer to BRAF inhibition. Mol Carcinog 55:1678–1687.

37. Wagner, J.P., Wolf-Yadlin, A., Sevecka, M., Grenier, J.K., Root, D.E., Lauffenburger, D.A., and MacBeath, G. 2013. Receptor tyrosine kinases fall into distinct classes based on their inferred signaling networks. Sci Signal 6:ra58.

38. Orian-Rousseau, V., Morrison, H., Matzke, A., Kastilan, T., Pace, G., Herrlich, P., and Ponta, H. 2007. Hepatocyte growth factor-induced Ras activation requires ERM proteins linked to both CD44v6 and F-actin. Mol Biol Cell 18:76–83.

39. Furge, K.A., Kiewlich, D., Le, P., Vo, M.N., Faure, M., Howlett, A.R., Lipson, K.E., Vande Woude, G.F., and Webb, C.P. 2001. Suppression of Ras-mediated tumorigenicity and metastasis through inhibition of the Met receptor tyrosine kinase. Proc Natl Acad Sci U S A 98:10722–10727.

40. Xu, Y.C., Wang, X., Chen, Y., Chen, S.M., Yang, X.Y., Sun, Y.M., Geng, M.Y., Ding, J., and Meng, L.H. 2017. Integration of Receptor Tyrosine Kinases Determines Sensitivity to PI3Kalpha-selective Inhibitors in Breast Cancer. Theranostics 7:974–986.

41. Raimondo, L., D’Amato, V., Servetto, A., Rosa, R., Marciano, R., Formisano, L., Di Mauro, C., Orsini, R.C., Cascetta, P., Ciciola, P., et al. 2016. Everolimus induces Met inactivation by disrupting the FKBP12/Met complex. Oncotarget 7:40073–40084.

42. Pfister, N.T., Fomin, V., Regunath, K., Zhou, J.Y., Zhou, W., Silwal-Pandit, L., Freed-Pastor, W.A., Laptenko, O., Neo, S.P., Bargonetti, J., et al. 2015. Mutant p53 cooperates with the SWI/SNF chromatin remodeling complex to regulate VEGFR2 in breast cancer cells. Genes Dev 29:1298–1315.

43. Li, D., Yallowitz, A., Ozog, L., and Marchenko, N. 2014. A gain-of-function mutant p53-HSF1 feed forward circuit governs adaptation of cancer cells to proteotoxic stress. Cell Death Dis 5:e1194.

44. Rahrmann, E.P., Moriarity, B.S., Otto, G.M., Watson, A.L., Choi, K., Collins, M.H., Wallace, M., Webber, B.R., Forster, C.L., Rizzardi, A.E., et al. 2014. Trp53 haploinsufficiency modifies EGFR-driven peripheral nerve sheath tumorigenesis. Am J Pathol 184:2082–2098.

